# Spatiotemporal Metabolic Responses to Water Deficit Stress in Distinct Leaf Cell-types of Poplar

**DOI:** 10.1101/2023.11.30.569476

**Authors:** Vimal Kumar Balasubramanian, Dusan Velickovic, Maria Del Mar Rubio Wilhelmi, Christopher R Anderton, C. Neal Stewart, Stephen DiFazio, Eduardo Blumwald, Amir H. Ahkami

## Abstract

The impact of water-deficit (WD) stress on plant metabolism has been predominantly studied at the whole tissue level. However, plant tissues are made of several distinct cell types with unique and differentiated functions, which limits whole tissue ‘omics’-based studies to determine only an averaged molecular signature arising from multiple cell types. Advancements in spatial omics technologies provide an opportunity to understand the molecular mechanisms underlying plant responses to WD stress at distinct cell-type levels. Here, we studied the spatiotemporal metabolic responses of two poplar leaf cell types-palisade and vascular cells-to WD stress using matrix-assisted laser desorption Ionization-mass spectrometry imaging (MALDI-MSI). We identified unique WD stress-mediated metabolic shifts in each leaf cell type when exposed to early and prolonged WD and recovery from stress. During stress, flavonoids and phenolic metabolites were exclusively accumulated in leaf palisade cells. However, vascular cells mainly accumulated sugars during stress and fatty acids during recovery conditions, highlighting a possibility of interconversion between sugars and fatty acids under stress and recovery conditions in vascular cells. By comparing our MALDI-MSI metabolic data with whole leaf tissue gas chromatography-mass spectrometry (GC-MS)-based metabolic profile, we identified only a few metabolites that showed a similar accumulation trend at both cell-type and whole leaf tissue levels. Overall, this work highlights the potential of the MSI approach to complement the whole tissue-based metabolomics techniques and provides a novel spatiotemporal understanding of plant metabolic responses to WD stress. This will help engineer specific metabolic pathways at a cellular level in strategic perennial trees like poplars to help withstand future aberrations in environmental conditions and to increase bioenergy sustainability.

## Introduction

Water-deficit (WD) stress is detrimental to plant growth and productivity. Plant responses to WD are dynamic and involve complex cross-talk between different regulatory pathways (Long, 2011), including metabolic adjustments and gene/protein expression at the molecular level for physiological and morphological adaptation at the whole-plant level (Dinneny et al., 2008). However, each cell-type in plant tissues is defined by specific transcriptional, protein, and metabolic profiles that determine its function and response(s) to stress (Plant Cell Atlas et al., 2021). With the establishment of the Plant Cell Atlas (PCA) community (Rhee et al., 2019), the importance of location-to-function paradigm has been highlighted, suggesting that this model has the potential to unlock new discoveries in plant sciences including how plants respond to environmental perturbations like WD (Plant Cell Atlas et al., 2021). Indeed, recent plant single-cell studies, which have been mostly performed in model species, indicate that plant responses to internal and external cues are largely cell type-specific (Xu and Jackson, 2023, Yang et al., 2021, Farmer et al., 2021, Nolan et al., 2023). Thus, determining the plant responses to WD requires the study of the cell/molecular properties of specific cell-types within a tissue to effectively reveal the underlying mechanisms regulating physiological processes and plasticity under suboptimal conditions.

Poplar (*Populus spp*.), a prevalent woody feedstock for improved next-generation biofuels, is one of the most biomass-productive native tree taxa in the northern hemisphere. While poplar species are good targets for carbon sequestration (Wullschleger et al., 2005) and production of feedstocks for biofuels and biomaterials (Bryant et al., 2020), they have also been widely used in physiological studies to understand how perennial trees respond to environmental changes (Li et al., 2014). Whole tissue responses to WD stress have been investigated in *Populus* at epigenome (Sow et al., 2021), transcriptome (Robertson et al., 2022, Rosso et al., 2023, Lee et al., 2021, Yang et al., 2023, Jia et al., 2017, Cossu et al., 2014, Wilkins et al., 2009), proteome (Plomion et al., 2006, Gao et al., 2022, Li et al., 2014, Xiao et al., 2009, Durand et al., 2011), and metabolome (Jia et al., 2020, Barchet et al., 2013, Hamanishi et al., 2015, He et al., 2022, Law, 2020, Tschaplinski et al., 2019b, Barchet et al., 2014) levels. However, spatiotemporal molecular mechanisms controlling plant responses to WD stress is not yet thoroughly understood in *Populus* and similar perennial trees.

Metabolites represent the downstream products of multiple interactions between genes, transcripts, and proteins. Small changes in gene and protein expression at functional levels are basically amplified at the metabolite level, which nominates metabolites as the ideal predictive molecular markers for stress response phenotypes compared to transcripts or proteins alone. Moreover, nonfunctional changes in many genes and proteins are not reflected in metabolites (Shen et al., 2023). Therefore, metabolomics has been considered as a very powerful approach for identifying the key primary and secondary small bioactive molecules with important roles in plant responses to environmental changes (Ahkami et al., 2019, Kumar et al., 2021, Barchet et al., 2013). The effect of WD stress at the metabolite level has been studied extensively in whole tissues using conventional extraction and metabolomics assays that include gas and liquid chromatography-based mass spectrometry (GC- and LC-MS) techniques (Jia et al., 2020, Hamanishi et al., 2015, Barchet et al., 2013, Nakayasu et al., 2016). Several studies in poplar reported whole leaf tissue level metabolomic shifts in response to WD stress conditions (He et al., 2022, Law, 2020, Tschaplinski et al., 2019b, Barchet et al., 2014). For instance, metabolomic analysis of whole leaf tissue of the WD-tolerant (*Populus simonii*) and -susceptible (*Populus deltoides* cv. ‘Danhong’) varieties suggested potential roles of increased antioxidants and long chain fatty acids in response to WD (Jia et al., 2020). Moreover, metabolomic analysis identified an increased level of raffinose family oligosaccharides in leaves of *P. balsamifera* under WD (Hamanishi et al., 2015). In another study with hybrid poplar (*P. deltoides* var. occidentalis and *P. × petrowskyana*), phenolic compounds, raffinose family-related compounds, and certain antioxidant metabolites were found to be involved in plant responses to WD stress (Barchet et al., 2013). However, our understanding of cell type specific metabolic changes under WD is scarce. Essentially, leaf cell types vary widely in their biological functions, in which palisade mesophyll cells are involved mainly in photosynthesis, while vascular cells are involved in inter-tissue signaling via xylem and phloem sap exchange. Therefore, cell-specific responses to WD at metabolic level may be critical for understanding their roles in plant stress tolerance.

Matrix-assisted laser desorption/ionization (MALDI) mass spectrometry imaging (MSI) is a robust molecular imaging technology that uses a focused laser beam to ablate and ionize material into the mass analyzer, which can provide high spatial resolution of the location of endogenous molecules from tissues *in situ*. Combined with spatial probing, a MALDI-MSI experiment enables simultaneous visualization of hundreds of molecules mapped to tissue morphology (Veličković et al., 2021). This approach has been used to evaluate the surface distribution of polysaccharides (Veličković et al., 2014), small sugars and glucosides (Bøgeskov Schmidt et al., 2018), metabolites, and lipids (Veličković et al., 2021) in different plant tissues and organs. For example, MALDI-MSI was used in barley roots to uncover the spatial distribution of metabolites in response to salinity stress (Sarabia et al., 2018). A recent advancement in spatial metabolic imaging approach using MALDI-MSI has been used to identify WD stress responses in *Piper sp.* and *Hibiscus rosa* sinensis root tissue (Honeker et al., 2022). The results showed a vascular and epidermis cell-specific accumulation of lignin-like metabolites, antioxidants, and fatty acid metabolites in each plant species, thereby highlighting the potential use of such a spatial imaging platform to visually map the metabolites in different cell types of WD-exposed plant tissues.

Here, we report spatiotemporal changes in metabolites levels in poplar leaf tissue when exposed to WD and recovery from stress. We describe the palisade specific accumulation of flavonoid and other phenolic compounds during WD treatment. In addition, accumulation of sugars together with the decreased number of fatty acids may indicate synthesis and transport of sugar via gluconeogenesis pathway during WD. Vascular cells showed an interconversion between fatty acids and sugar levels during WD and recovery period. Overall, our results unravel cell-specific spatiotemporal metabolic changes in response to WD. Further, we report the use of MALDI-MSI as a powerful technique when combined with conventional GC-MS-based whole tissue metabolic profiling to gain a granular understanding of plant responses to WD stress. This work highlights the power of a spatial metabolomics approach in understanding plant responses to abiotic stresses, enabling future attempts at mapping molecular machineries to cellular domains. These mechanistic molecular insights can be exploited to aid the design of strategic bioenergy trees with enhanced tolerance to WD stress.

## Materials and methods

### Plant growth conditions and water-deficit stress treatment

Clones of *Populus tremula* x *alba* (INRA 717 1-B4) were rooted in sterile conditions in a growth chamber (16-hr/8-hr day/night and 24°C/18°C 600 μmol m^−2^s^−1^) for at least 25 days. Seedlings were transplanted into 1.5 L pots filled with Profile Porous Ceramic (PPC) soil. The seedlings were grown (16-hr/8-hr day/night and 24°C/18°C 800 μmol m^−2^s^−1^) in the greenhouse and fertilized every day with a nutrient solution containing N 77 ppm, P 20 ppm, K 75 ppm, Ca 27 ppm, Mg 17 ppm, S 65 ppm, Fe 1.50 ppm, Mn 0.50 ppm, Zn 0.05 ppm, Mo 0.01 ppm, Cu 0.02 ppm and pH 5.6. Water deficit treatments were applied 45 days after rooting by withholding water until visual stress symptoms (i.e., leaf wilting) appeared (30–35% relative soil water content) (early water deficit, E-WD). Plants were kept for ten days at 35% soil water content (late water deficit, L-WD) and then re-watered with regular fertilization for three days after sampling for the recovery period (REC). Samples from each treatment were collected between 9-10 am (1 and ½ –2 and ½ hours after the light turned on). Midrib-containing leaf punches were harvested, flash frozen in liquid nitrogen, and kept at −80°C until use. These leaf punches (disks with ∼0.5-inch diameter in size) were used for MALDI-MSI workflow for spatial metabolomic analysis. Alternatively, whole intermediate leaves from an independent WD experiment were harvested, immediately frozen in liquid nitrogen, and kept at −80°C until use. These whole leaf tissues were used for GC-MS-based metabolomic profiling. At any given stress or recovery timepoints, leaf punches were collected from six biological replicates for MALDI-MSI, while the entire leaf from three independent biological replicates were collected for whole leaf metabolite profiling. Leaf gas exchange parameters were collected from four biological replicates.

### Gas-exchange and biomass measurements

Photosynthesis measurements were recorded in intact plants using a portable gas exchange system (LI-COR 6400). Photosynthesis was induced by saturating light (1000 μmol m^−2^ s^−1^) with 400 μmol mol^−1^ CO_2_ surrounding the leaf (Ca). The amount of blue light was set to 10% photosynthetically active photon flux density to optimize stomatal aperture. Block temperature was set to 24 °C. Plant samples for biomass assay were collected and oven dry at 60 °C for 5 days.

### Cryosectioning of frozen leaf tissue and sample preparation for MALDI-MSI imaging

Leaves were embedded within a mixture of 7.5% hydroxypropyl methylcellulose (HPMC) and 2.5% polyvinylpyrrolidone (PVP), and 12 µm sections were thaw mounted on indium tin oxide (ITO)-coated glass slides using a cryotome maintained at -14°C (CryoStar NX-70 Cryostat, Thermo Scientific, Runcorn, UK). Slides were vacuum dried and homogenously sprayed, using a M5 Sprayer (HTX Technologies, Chapel Hill, NC), with 2,5-dihydroxybenzoic acid (DHB) matrix for analysis in positive ion mode(Veličković et al., 2021) and N-(1-naphthyl) ethylenediamine dihydrochloride (NEDC) matrix for analysis in negative ion mode(Honeker et al., 2022). DHB was prepared at a concentration of 40 mg/mL DHB (in 70% MeOH) and was sprayed at 50 µL/min flow rate. The nozzle temperature was set to 70 °C, with 12 cycles at 3 mm track spacing with a crisscross pattern. A 2 s drying period was added between cycles, a linear flow was set to 1,200 mm/min with 10 PSI of nitrogen gas and a 40 mm nozzle height. This resulted in matrix coverage of ∼667 µg/cm^2^ for DHB. NEDC was prepared at a concentration of 7 mg/mL NEDC in 70% MeOH and was sprayed at 120 µL/min flow rate. The nozzle temperature was set to 70 °C, with 8 cycles at 3 mm track spacing with a crisscross pattern. A 0 s drying period was added between cycles, a linear flow was set to 1200 mm/min with 10 PSI of nitrogen gas and a 40 mm nozzle height. This resulted in matrix coverage of ∼187 µg/cm^2^ for NEDC.

### MALDI-MSI data collection and processing

All imaging analyses were performed on an ESI/MALDI dual-source MALDI Fourier transform ion cyclotron resonance (FTICR)-MS equipped with the ParaCell (scimaX 2XR 7T, Bruker Daltonics, Bremen, Germany) operating in MALDI mode with the data point sizes of 2 M and 2w detection. Positive ion mode acquisitions with DHB were acquired with broadband excitation from *m/z* 92 to 1,000, resulting in a detected transient of 0.418 s— the observed mass resolution was ∼200k at *m/z* 400. Negative ion mode NEDC analyses were acquired with broadband excitation from *m/z* 92 to 1,000, resulting in a detected transient of 0.418 s— the observed mass resolution was ∼200k at *m/z* 400. FlexImaging (Bruker Daltonics, v.5.0) was used for the imaging acquisition, and analyses were performed with 30 µm step size. FlexImaging sequences were directly imported into SCiLS Lab (Bruker Daltonics, v.2023.a Premium 3D) using automatic MRMS settings. Ion images were directly processed from the profile datasets within SCiLS Lab, and automated annotation of the centroided dataset was completed within METASPACE. KEGG-v1 and SwissLipids were used as databases for annotaions within 3 ppm m/z tolerance. List of annotated m/z features was imported back to the SCiLS, and table with ion intensity of each annotated feature (after RMS normalization) in palisade and vascular part of each leaf section was created and used further for statistical (t-test) analysis. Palisade and vascular region of the leaf were manually outlined in the SCiLS software (Bruker Daltonics) using brightfield microscopy image recorded for each analyzed section. Due to instrument and resource limitations, we were not able to collect leaf punches from the entire leaf for sectioning and MALDI-MSI. However, we used 6 biological replicates with 2 technical replicates (each consecutive sections, n=12) for any given condition. The relative abundance level of a metabolite in palisade or vascular cell was calculated by summing the relevant abundance levels from a region of tissue comprising cell-types of each kind (Supplementary Figure S2).

### Whole leaf tissue metabolites profiling by GC-MS

Ground frozen powder of the intermediates leaves from poplar plants grown under control, and water deficit (E-WD and L-WD) and recovery conditions were submitted to the West Coast Metabolomics Center (University of California, Davis), extracted, measured, and analyzed by gas chromatography–mass spectrometry (MS) (Gerstel CIS4–with a dual MPS Injector/Agilent 6,890 GC-Pegasus III TOF MS) as described before (Weckwerth, Wenzel, & Fiehn, 2004). Processes for the integrated extraction, identification, and quantification of metabolites were performed according to Fiehn et al. (2008). Metabolites are expressed as differential abundancy of normalized intensity values (log2) between Control and E-WD, L-WD and recovery treatments.

### Van-Krevelen classification and pathway analysis

Van-Krevelen analysis was performed as reported (Honeker et al., 2022). Briefly the molecular formula from significantly upregulated and downregulated metabolites (t-test, p<0.05) was used to classify them into higher order metabolic classes such as lignins, hydrocarbons, tannins, carbohydrates, proteins, etc., based on the elemental ratios O/C and H/C ratios. Molecular formulas that come under broader classes, such as lignins, were looked up in KEGG pathways to identify specific sub-class, such as flavonoids, to derive pathway maps in Figures 2 and 3.

### Statistical analysis

For biomass, leaf gas exchange parameters, and MALDI-MSI relative metabolite abundance analysis, the significance test was performed using T-tests with the built-in statistical function in Microsoft Excel. For the whole leaf tissue metabolomics analysis (GC-MS), the significance test was performed using one-way ANOVA analysis. ANOVAs of individual features were performed using aov function of STATS package in R (R Core Team 2021). In this manuscript, we present significantly changed data under stress or recovery conditions (P < 0.05) compared with controls; however, in the case of MALDI-MSI analysis, we also include the moderately changed data (0.5 < P ≤ 0.1) compared with controls.

## Results

### Plant biomass and photosynthesis rates were affected by water deficit stress

Whole plant responses to water limitation were monitored via biomass and gas exchange parameter measurements during early WD (E-WD), late WD (L-WD), and recovery (R) conditions. Compared to the control well-watered condition, plant shoot biomass was reduced under E-WD (-48%), L-WD (-67%) and R (-75%) treatments (Figure S1A, B; Supplementary table S1). Root biomass was reduced during E-WD (-44%) and R (-65%) but showed no change during L-WD compared to control roots (Fig S1B). Among physiological parameters, leaf tissue showed a decrease in net photosynthesis (A_N_) (-75%, -23%), stomatal conductance (g_s_) (-95%, -61%), and leaf transpiration (-91%, -51%) during E-WD and L-WD stages, respectively. In contrast, during recovery from stress, higher values of g_s_ (+80%) and leaf transpiration (+58%) were observed compared to non-stressed control plants (Supplementary Figure S1B and Supplementary table S1).

### A spatiotemporal metabolic shift was detected in poplar leaf cell types

Leaf tissues were harvested from plants under WD and recovery conditions, cryosectioned, and used for MALDI-MS imaging-based spatial metabolomics. We looked for metabolites that were significantly altered by WD treatment compared to controls, especially in palisade mesophyll and vascular regions of cryosectioned leaf tissue (Supplementary Figure S2). By combining positive and negative ionization mode analyses in MALDI, a total of 2,358 spectral features were detected, which were then curated to remove false positives and spotty non-real abundance patterns. This left 103 molecular features that were significantly altered in palisade and/or vascular cell types under E-WD/L-WD and/or recovery time points compared to the control well-watered (WW) condition (Supplementary Table S2). Of those, 60 molecular features were significantly altered in a cell type-specific manner (strictly enriched or depleted in palisade or vascular cell type), while 43 metabolites were significantly altered in both leaf cell types. Between time points, recovery condition had more metabolites altered in both palisade (n=29) and vascular (n=57) cell types compared to E-WD and L-WD conditions (Figure 1A). However, between the two cell types, vascular cells showed a higher number of significantly altered metabolites than palisade cells under any harvesting timepoints (Figure 1A). We categorized the identified molecular features into metabolic classes through Van Krevelen classification (Honeker et al., 2022), which groups molecular formulas based on their elemental ratios (O/C and H/C ratios) and degree of unsaturation. The categories included different biochemical classes such as secondary metabolites (lignin, condensed hydrocarbons, unsaturated hydrocarbons), lipids, and carbohydrate-related metabolites (Figure 1B, Supplementary Table S2). This analysis showed a higher number of WD- or recovery-altered metabolites belonged to lignins (n=39) and condensed hydrocarbons (n=33), followed by carbohydrates (n=16), lipids (n=3), and other metabolic classes (n=8) (Figure 1B). The MALDI-MSI technique used in this study cannot resolve between stereoisomers, which means that the relative abundance levels of a molecular feature represent an average abundance of all stereoisomers (e.g., C_6_H_12_O_6_ isomers include monosaccharides such as glucose, fructose, mannose, etc.). Therefore, any significant changes in the cellular abundance level of C_6_H_12_O_6_ represent a significant shift in total monosaccharide levels in a particular cell type. Based on the metabolite classification outcome (Figure 1B) and to further identify distinct spatial trends in leaf cell types, we mainly focused on secondary metabolites, lipids, and carbohydrate-related metabolites in two major leaf cell types, i.e., palisade mesophyll and vascular cell types, when exposed to WD conditions. We identified a palisade cell type specific metabolic trend in flavonoid biosynthesis pathway, while in vascular cells, lipid and sugar metabolism altered in a cell type specific manner.

**Figure 1.**
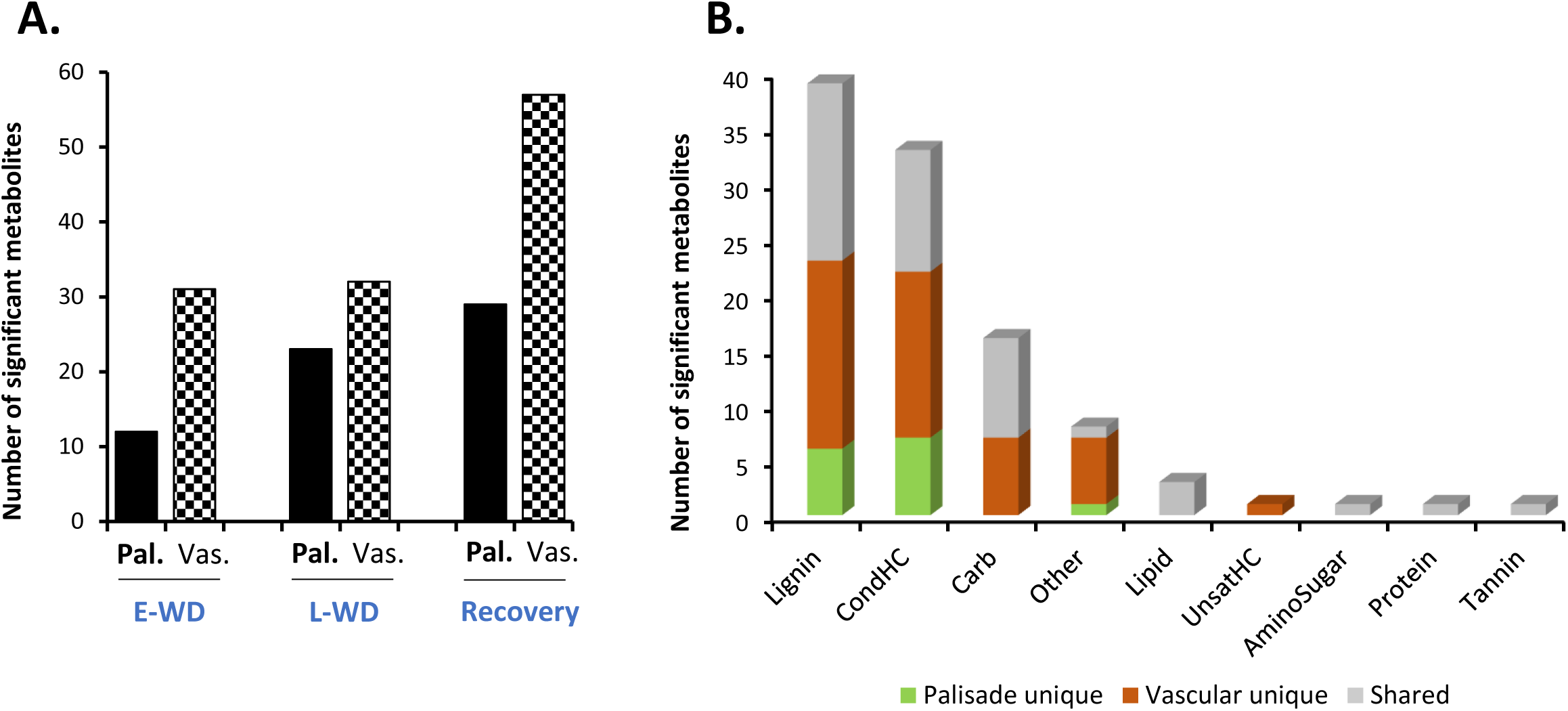
Spatial metabolomics identified cell type-specific WD stress-responsive metabolites in Poplar leaf. **(A).** Total number of significantly altered metabolites under early water deficit (E-WD), late water deficit (L-WD), and recovery conditions identified through MALDI-MSI analysis. Metabolites significantly altered in two major leaf cell types, palisade (Pal.) and vascular (Vas.) cells, are marked with different bar patterns. The relative abundance level of each significant metabolite was averaged from 6 biological replicates with 2 technical replicates in any given condition. T-test was used for statistical analysis. Metabolites with significant (n=97, p<0.05) and moderate significant (n=7, 0.05<p≤0.1) changes are included. **(B).** Metabolites were grouped into various metabolic classes by Van Krevelen classification and cell-type specificity of metabolites under each metabolic class were highlighted with green (Pal unique), brown (Vas unique) and grey (Pal and Vas shared metabolites) colors.

### Water deficit stress elevated the levels of specific flavonoids and fatty acids uniquely in leaf palisade cells

Many plant species, including poplars, were shown to elevate the levels of different flavonoids in whole leaf tissues under water-limited conditions that help scavenge the ROS generated during stress(Ahmed et al., 2021b, Shomali et al., 2022, Tattini et al., 2004). Here, our spatial metabolomics approach allowed us to look for any cell type level differences in flavonoid distribution during WD stress and recovery time points in hybrid poplar leaf. Palisade mesophyll cells uniquely exhibited a higher relative abundance of specific flavonoids during E-WD, L-WD, and recovery stages (Figure 2A and Supplementary Table S2). Compared to the E-WD stage, prolonged WD stress induced the accumulation of a greater number of flavonoids in a cell-type specific manner. Among those, tremulacin was increased (+2.89-fold) in palisade cells during E-WD, while stereoisomers quercetin and tricetin, represented by the molecular formula C_15_H_10_O_7_, were elevated (+1.45-fold) exclusively during L-WD stage. Prolonged WD also induced the abundance levels of C_9_H_8_O_4_ that represent caffeate (+2.4-fold), and polyketides such as pseudopurpurin (+1.33-fold) exclusively in palisade cells (Figures 2A, 2B and Supplementary Table S2). During recovery from WD stress, a higher level of C_15_H_10_O_6,_ which represents stereo-isomeric flavonoids such as kaempferol and luteolin (+1.24-fold), C_14_H_8_O_5_ which include purpurin (+1.28-fold), and two aromatic phytochemicals riccionidin A (+1.33-fold) and rhein (+1.37-fold) were exclusively detected in palisade cells (Figures 2A, 2B and Supplementary Table S2). Vascular cells did not show any significant changes in abundance levels of the abovementioned flavonoids during the WD or R stages (Figures 2A, 2B and Supplementary Table S2). Therefore, palisade cell-specific accumulation of these secondary metabolites, mostly during L-WD and recovery stages, highlights a spatially distinct metabolic activity in palisade cells in response to WD stress and recovery in poplar. However, a number of other secondary metabolites that accumulated in vascular cells had reduced abundance in palisade cells. They include C_13_H_18_O_7_, C_7_H_14_N_2_O_7_, and C_7_H_11_NO_7_P_2,_ representing salicin (+1.62-fold), coumeroic acid (+2.24-fold), and risedronic acid (+1.71-fold), respectively (Supplementary Table S2).

**Figure 2.**
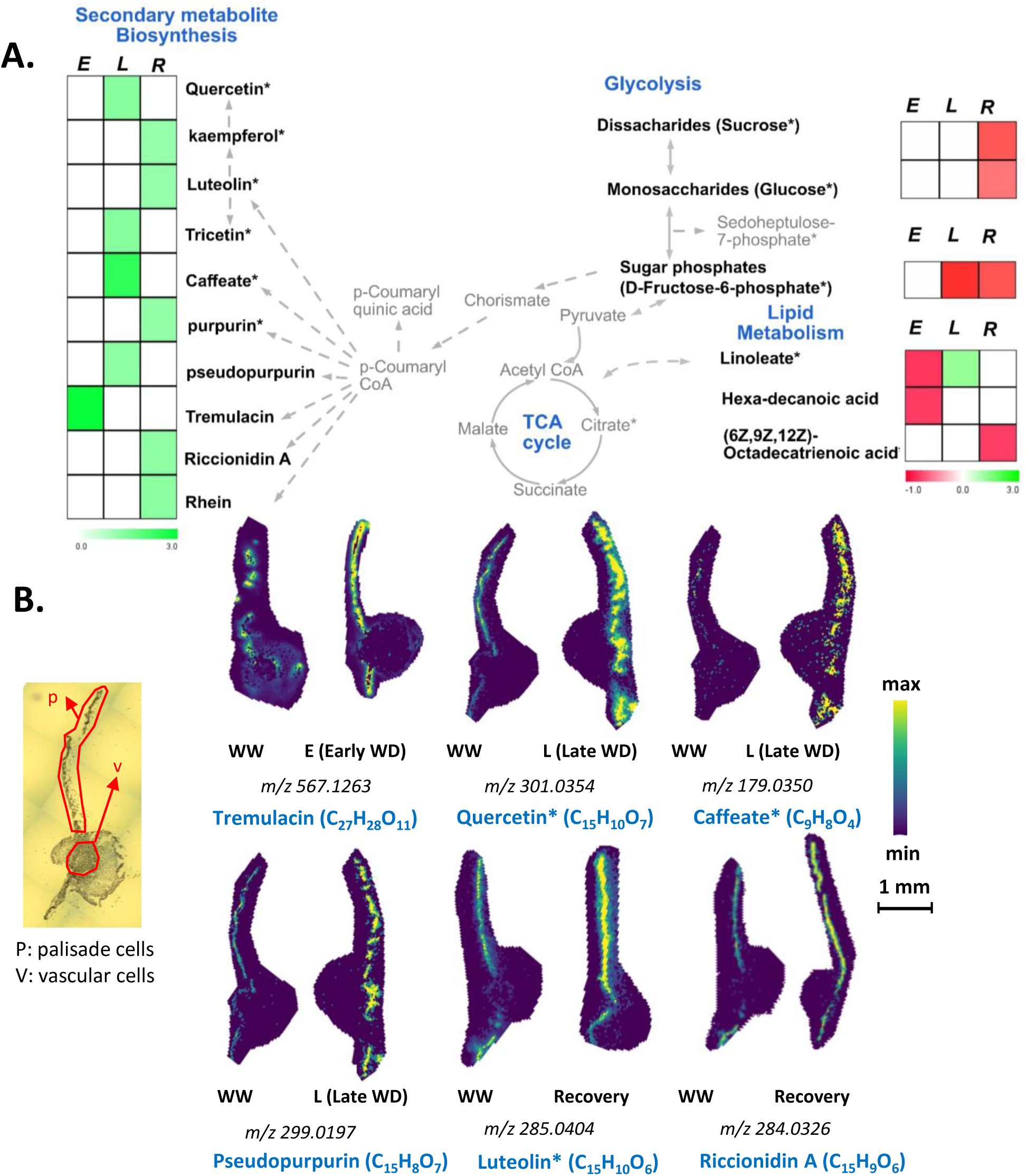
Palisade cell type-specific metabolites identified by MALDI-MSI analysis. **(A).** Metabolic pathway map of selected palisade cell-type abundant metabolites that were significantly altered during early (E), late (L) water-deficit stress and recovery (R) conditions. Metabolites with significant increased and decreased abundance levels during stress or recovery conditions compared to the control well-watered condition are presented by green and red colors, respectively. Metabolite relative abundance levels were averaged from 6 biological replicates with 2 technical replicates in any given condition. T-test was used for statistical analysis and metabolites that were significant (p<0.05) and moderately significant (0.05<p ≤ 0.1) were used to generate heatmaps. Solid and dashed arrows represent direct and indirect metabolic pathway relationships, respectively. **(B).** Selective ion images of metabolites with palisade cell-type unique enrichment are shown. The ion image for each metabolite is a representative figure from 6 biological replicates with two technical replicates. Metabolite names with an asterisk (*) contain potential stereoisomers sharing the same molecular formula. For example, Quercetin* (C_15_H_10_O_7_) and caffeate* (C_9_H_8_O_4_) are shown as representative metabolites among the stereoisomers with same molecular formula. The scale bar is 1mm. WW: well-watered condition; WD: water-deficit stress.

Unsaturated fatty acids also play a role in stress response by scavenging ROS (He and Ding, 2020b). During L-WD stress, we noticed a moderate increase in C_18_H_32_O_2_ representing the fatty acid linoleate (+1.22-fold) only in palisade cells (Figure 2A), while its abundance level did not change in vascular cells, highlighting a potential role for higher linoleate levels in palisade cells during L-WD stress. Moreover, two other fatty acids, hexadecanoic acid (-0.75-fold) and (6Z,9Z,12Z)-octadecatrienoic acid (-0.78-fold), showed reduced levels in palisade cells during the E-WD and R timepoints (Figure 2A, Supplementary Table S2). Altogether, the WD-stressed palisade cells showed a cell-type specific accumulation of flavonoids and the fatty acid linoleate, especially under L-WD and recovery stages, which was not observed in the vascular cell-type (Figures 2A, 2B and Supplementary table S1).

### Vascular cells showed an opposite trend with the accumulation of fatty acids and sugars in response to water deficit stress

Vascular cells transport a wide array of primary and secondary metabolites via xylem and phloem sap and provide structural support to the plant through secondary cell wall formation. Here, using a spatial metabolomics approach, we found an opposite trend with the levels of fatty acids and glycolytic metabolites in vascular cells especially during E-WD and Recovery conditions (Figure 3A, 3B). During E-WD stage, vascular cells showed reduced levels of three fatty acids, linoleate (-0.73-fold), hexadecanoic acid or palmitic acid (-0.84-fold), and C_18_H_30_O_2_ that potentially represent (6Z,9Z,12Z)-Octadecatrienoic acid (-0.73-fold), a polyunsaturated fatty acid (Figure 3A, 3B & Supplementary Table S2). However, focusing on the glycolysis/gluconeogenesis pathways, we observed an increase in sugars and sugar-phosphate levels under E-WD condition (Figure 3A,3B). The level of C_12_H_22_O_11,_ which comprises disaccharide sugars such as sucrose and maltose showed a +1.34-fold increase, while C_6_H_12_O_6,_ which comprise monosaccharide sugars such as glucose, fructose, and mannose, showed a +2-fold increase exclusively in vascular cells during E-WD condition (Figure 3A, 3B). Although these di- and mono-saccharides were also detected in palisade cells (Figure 2A), their abundance levels were not significantly altered under E-WD stress vs. control conditions. Moreover, sedoheptulose 7-phosphate and related isomers (C_7_H_15_O_10_P) was also highly abundant in vascular cells (+1.56-fold) under E-WD stress condition. Under recovery condition, however, we found an opposite trend with the abundance levels of the abovementioned sugars and fatty acids, in a way that the levels of disaccharides (-0.63 fold) and monosaccharides (-0.69 fold) were reduced, while the relative abundance levels of all three fatty acid levels including linoleate (+1.48 fold), hexadecanoic acid (+1.18-fold), and (6Z,9Z,12Z)-octadecatrienoic acid (+1.74 fold) were increased. The observed opposite trend between fatty acid, mono, and disaccharide sugars, and sugar-phosphate levels during stress and recovery conditions indicate that vascular cells possibly break down the fatty acids to generate sugars via the gluconeogenesis pathway to support respiration during WD stress conditions. Moreover, the abundance level of C_6_H_8_O_7,_ which represents the TCA cycle intermediate citrate and its isomers, was increased during recovery (+1.58 fold), highlighting a possible role of citrate in respiration and activation of fatty acid biosynthesis when plants are recovered from WD (Pracharoenwattana et al., 2005). The other TCA cycle-related metabolites, such as cis-aconitate and oxoglutarate, showed no significant changes in their abundance levels in stress and recovery stages (supplementary table S2).

**Figure 3.**
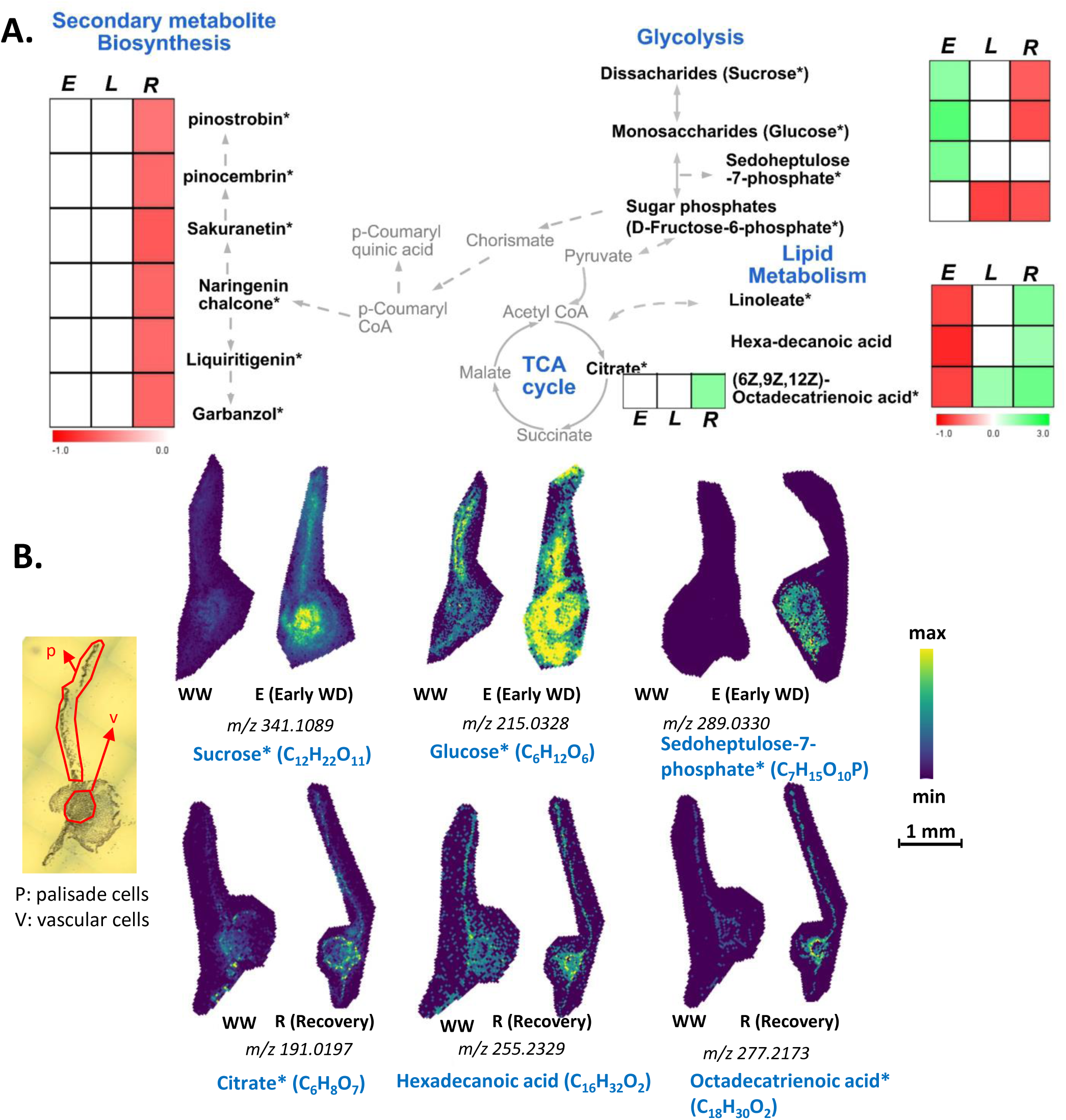
Vascular cell type-specific metabolites identified by MALDI-MSI analysis. **(A).** Metabolic pathway map of selected vascular cell-type abundant metabolites that were significantly altered during early (E), late (L) water-deficit stress and recovery (R) conditions. Metabolites with significant increased and decreased abundance levels during stress or recovery conditions compared to the control well-watered condition are presented by green and red colors, respectively. Metabolite relative abundance levels were averaged from 6 biological replicates with 2 technical replicates in any given condition. T-test was used for statistical analysis and metabolites that were significant (p<0.05) and moderately significant (0.05<p ≤ 0.1) were used to generate heatmaps. Solid and dashed arrows represent direct and indirect metabolic pathway relationships, respectively. **(B)**. Selective ion images of metabolites with vascular cell-type specific enrichment are shown. The ion image for each metabolite is a representative figure from 6 biological replicates with two technical replicates. Metabolite names with an asterisk (*) contain potential stereoisomers sharing the same molecular formula. For example, sucrose* (C_12_H_22_O_11_) and Glucose* (C_6_H_12_O_6_) are shown as representatives of disaccharides and monosaccharide stereoisomers, respectively. The scale bar is 1mm. WW: well-watered condition; WD: water-deficit stress.

Apart from fatty acids and sugar metabolites, vascular cells showed a reduced level of 27 different secondary metabolites during recovery from WD stress (Figure 3A and Supplementary Table S2). These include C_15_H_12_O_5,_ which represents flavonoid naringenin chalcone (-0.58-fold), and several other phenylpropanoid pathway-derived metabolites such as dihydrokaempferol (-0.56-fold), 4’,6-Dihydroxyflavone (-0.52-fold), and 4’-O-Methylisoflavone (-0.50-fold). (Supplementary table S2). Such an overall reduction in secondary metabolite biosynthesis in vascular cells during recovery from stress indicates a possible spatiotemporal diversion of carbon flow to other essential primary metabolism pathways to support recovery from stress.

### MALDI-mass spectrometry imaging as a complementary technique to traditional whole tissue metabolite profiling

Whole tissue metabolomics approaches using GC-MS and LC-MS techniques have been widely used to address different questions in plant biology including plant responses to stress (Jorge et al., 2016, Ma and Qi, 2021). However, the outcomes of these studies have not been correlated with spatial and temporal distribution of metabolites in plant cell-types. Here, we performed a whole leaf tissue GC-MS-based metabolite profiling during E-WD, L-WD and recovery stages, and then compared the outcomes with our MALDI-MSI data. We hypothesized that spatial metabolomics verifies the whole tissue metabolic trends under WD condition and more importantly provides additional and key complementary information that are missed with traditional whole tissue-based approach. Out of 427 GC-MS-based metabolic features that changed significantly during stress and recovery conditions, 147 metabolites were identifiable through a library search. Among them, E-WD, L-WD, and recovery treatments resulted in 93, 74, and 55 significant metabolites, respectively (Supplementary Table S3, Supplementary Figure S3). We found 26 metabolites that were commonly detected by both GC-MS and MALDI approaches. However, compared to our spatial MALDI-MSI data, only seven metabolites showed a similar trend (increase or decrease) in their relative abundance levels in whole leaf tissue. They include glucose and mannose (increased during E-WD in whole leaf tissue) which regarding MALDI-MSI refer to monosaccharides (C_6_H_12_O_6_) (increased in vascular cells during E-WD) (Figure 4A, 4B); palmitate or hexadecanoic acid (increased under recovery condition in whole leaf tissue) which regarding MALDI-MSI refers to C_16_H_32_O_2_, (increased in vascular cells during recovery condition); and inositol-4-monophosphate, glucose-6-phosphate, galactose-6-phosphate, and fructose-6-phosphate (reduced during L-WD in whole leaf tissue) which regarding MALDI-MSI collectively refer to sugar phosphates (C_6_H_13_O_9_P) (reduced in both palisade and vascular cells during L-WD) (Figure 4A, 4B). This highlights the use of the MALDI-MSI as a powerful complementary tool to understand the spatial distribution of metabolites in plant tissues during environmental stresses.

**Figure 4.**
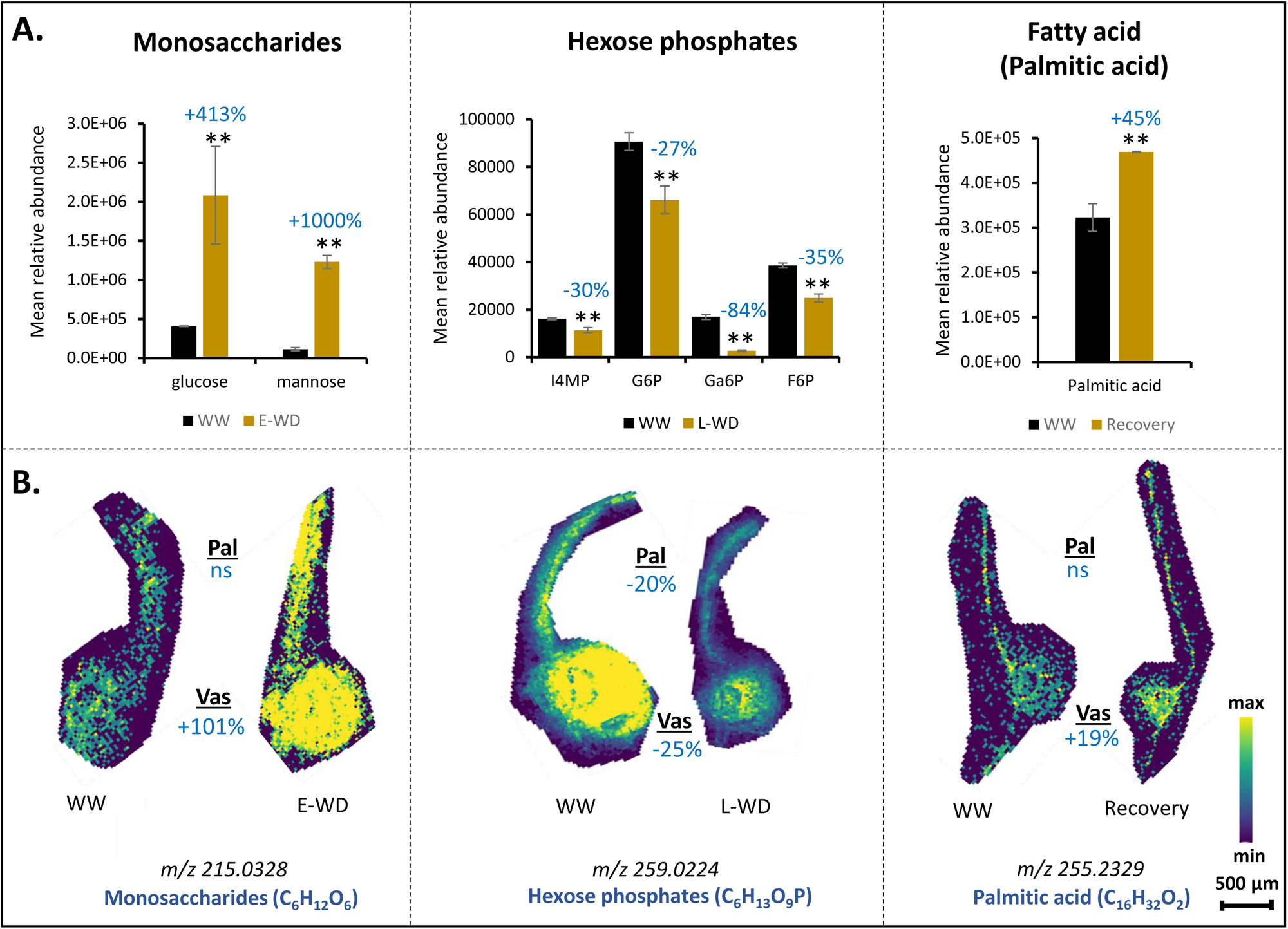
Metabolite accumulations at whole leaf tissue level corresponds to spatially resolved metabolites at cellular levels. **(A).** Relative abundance levels of selective metabolites identified by whole leaf tissue-based GC-MS metabolomics analysis under early water deficit (E-WD), late water deficit (L-WD), and recovery conditions. Metabolite abundance is calculated from an average of 3 biological replicates. One-way ANOVA was used for statistical analysis (** represents p<0.05). **(B).** Selective ion images of monosaccharide, sugar phosphates, and fatty acids that show similar accumulation patterns with whole leaf metabolite profiles during stress or recovery conditions are provided. For monosaccharides and hexose phosphates, the ion abundance image reflects all stereoisomers with similar molecular formula. Ion image for each metabolite is a representative figure from 6 biological replicate with 2 technical replicates. T-test was used for statistical analysis (0.1≤p<0.05). Levels of metabolic shift of each compound in stressed vs. control samples are shown by percentages in dark blue color. Pal: palisade; Vas: vascular; ns: not significant; WW: well-watered condition; WD: water-deficit stress.

## Discussion

Recently, several studies highlighted the importance of understanding plant cell type-specific abiotic stress responses to maximize plant performance under suboptimal environmental conditions (Jeong et al., 2013, Fàbregas et al., 2018, Lee et al., 2016). In this work, we studied the metabolic responses of poplar to WD in a spatiotemporal manner and identified several distinct metabolites or pathways uniquely activated in leaf tissue cell types.

Leaf development and patterning involve co-ordination between meristematic cell division, cell expansion, and differentiation processes that form specialized cell-types. The adaxial side of the leaf tissue is packed with chloroplast-dense palisade mesophyll cells which are involved in photosynthesis. In contrast, the vascular cells mobilize the fixed photosynthate and other macro molecules to other tissues. Several recent single-cell transcriptomic studies performed in leaf tissue of model plants highlighted the distinct molecular profiles associated with different leaf cell types (Tenorio Berrío et al., 2021, Zang et al., 2023, Zhu et al., 2023), re-emphasizing that palisade and vascular cells are developmentally and functionally distinct, possibly exhibiting cell-type unique responses to environmental stresses such as WD.

During abiotic stresses such as WD, cellular ROS levels increase, which leads to oxidative damage of many cellular processes, including fatty acid peroxidation (Cruz de Carvalho, 2008). Previous metabolomic analysis at the whole tissue level suggested important roles for flavonoids and aromatic compounds in drought stress tolerance in several plant species (Ahmed et al., 2021a, Shomali et al., 2022, Tattini et al., 2004). Using spatial metabolomics, we found an exclusive increase in the abundance of several secondary metabolites including flavonoids, polyketides, and phenylpropanoid pathway-related metabolites during L-WD stage in palisade cells (Figure 2A, 2B, Supplementary table S2). Among them, quercetin and luteolin are known to regulate ROS levels in plants by playing antioxidant roles (Singh et al., 2021, Hodaei et al., 2018, Gori et al., 2021). Other such as Riccionidin was found to confer abiotic stress tolerance in liverwort (*Marchantia polymorpha*), while anthraquinone derivatives such as purpurin and pseudopurpurin have been found to have antioxidant properties (Albert et al., 2018, Jin et al., 2011). The increased induction of secondary metabolism under prolonged drought in *Populus* was associated with adaptive mechanisms rather than metabolic perturbations, leading to increased organic solute accumulation (reduced osmotic potential) for improved drought tolerance (Tschaplinski et al., 2019a). Moreover, our results showed the accumulation of other aromatic compounds that remained undetected in previous studies where whole tissue metabolomics-based approaches were applied. For example, we found an increased level of tremulacin, a salicylate derivative (Feistel et al., 2015, Rubert-Nason et al., 2014), in a palisade-specific manner under E-WD condition. This highlights the role of palisade cells in exclusively directing carbon partitioning to secondary metabolism involved in ROS-neutralization and antioxidant responses to WD stress. Focusing on fatty acids, the levels of linoleate were increased in the palisade cells during the L-WD treatment, while no significant changes were observed in vascular cells (Supplementary Table S2). By anatomy, palisade mesophyll cells have a relatively high chloroplast density whose organellar membranes are composed mainly of unsaturated fatty acids (Hernández and Cejudo, 2021). Linoleate is an unsaturated fatty acid and is part of the chloroplast membrane. An increase in the degree of unsaturation of membrane lipids was associated with improved drought resistance and ROS neutralization (He and Ding, 2020a, Ullah et al., 2022, Zi et al., 2022), which highlights the potential role of fatty acid composition in poplar leaf palisade cells to help mitigate the L-WD stress-related consequences. Looking at leaf physiological parameters during WD stress (Supplementary Figure S1B), compared to E-WD, photosynthesis, leaf conductance, and transpiration rates were improved during L-WD, signifying the possible roles of flavonoids and fatty acids in neutralizing ROS (Jia et al., 2021, Resmi et al., 2015), leading to the observed improved leaf performance during L-WD stress.

Vascular cells establish inter-tissue signaling via xylem and phloem sap, through which essential metabolite classes, such as hormones, fatty acids, and sugars, are transported. Phloem cells were shown to transport various fatty acids during normal and stress conditions (Guelette et al., 2012, Madey et al., 2002). In our spatial MALDI-MS data, all three detected fatty acids (linoleate, hexadecanoic acid or palmitic acid, and (6Z,9Z,12Z)-octadecatrienoic acid) were decreased in abundance in vascular cells during E-WD stress conditions (Figure 3A, 3B, Supplementary Table S2). These results may indicate the degradation of these fatty acids, via beta oxidation for energy production, as WD stress generally reduces the photosynthetic carbon fixation capacity (Supplementary Figures S1). Activation of beta oxidation may be associated to the increased levels of disaccharide and mono-saccharides and pentose phosphate pathway-related metabolites during E-WD, possibly via gluconeogenesis pathway (Figure 3A, 3B). Moreover, during recovery, our time-course MALDI-MSI analysis showed an opposite trend between fatty acids and sugar abundance levels (Figure 3A, 3B), where we found an increased level of all three fatty acids and decreased accumulation of sugars exclusively in vascular cells. This vascular cell-specific opposite trend between fatty acid and sugar levels during WD stress and recovery conditions indicates that vascular cells could possibly break down fatty acids to generate sugars during WD stress and re-synthesize fatty acids during recovery (possibly by utilizing those sugars) as the demand for lipids is increased during recovery from stress (Gigon et al., 2004). In addition, during recovery from stress, a lower abundance was detected for many of the secondary metabolites exclusively in vascular cells (Supplementary Table S2), highlighting the possibility that vascular cells diverted the carbon flow from secondary metabolite biosynthesis to other essential pathways, such as fatty acid biosynthesis during WD stress.

Water deficit stress imposes severe phenotypic damage to annual plants and perennial trees such as poplars. Annual plants escape from drought stress by allocating carbon to seed production, thereby advancing to the next generation. However, due to a longer reproductive cycles, perennial trees must withstand stress through different strategies such as tolerance or avoidance mechanisms involving stomatal regulation, synthesizing osmoprotectants, including ROS-neutralizing metabolites. Although more studies have been conducted on responses of annual plants to WD, there is a lack of understanding of metabolic responses of perennial trees to WD. In our work, we observed an overall reduction in poplar aboveground biomass during both early and late stress stages, while an increase in root biomass was observed in L-WD (compared to E-WD) (Supplementary Figure S1B). This could possibly be associated with the observed higher sugar levels in leaf-vascular cells during E-WD and their possible translocation to roots to reserve carbon for later usage or for root growth in response to WD stress. Carbon mobilization and storage in belowground sinks such as roots during dormancy followed by re-mobilization of carbon to support bud outgrowth in spring has been commonly reported in trees. On the other hand, in oak saplings (Heizmann et al., 2001), it was reported that under reduced photosynthetic condition, a higher-level carbon was translocated from roots to leaves via xylem sap. Our hypothesis on vascular cells-mediated sugar transport to support root growth or from root system to support shoot growth and development especially during WD should be further tested by isotope labeling and root metabolite profiling in future work. Stress memory is another emerging area where plants accumulate stress-responsive molecular signatures, enabling them to withstand future stress events (Tombesi et al., 2018). In our study, the recovery from WD was found to have higher levels of flavonoids in palisades and higher levels of fatty acids in vascular cells, possibly highlighting a preparedness for future episodes of WD stress.

We employed a GC-MS based approach to profile the whole leaf tissue metabolic changes under WD stress and recovery conditions, and then compared the outcome with the spatial distribution of metabolites derived by MALDI-MSI. However, we found only a few metabolites that showed similar accumulation patterns between the spatial and whole tissue datasets (Figure 4A, 4B). This could be attributed to the inherent ionization, mechanical, and molecular selectiveness differences between these two mass spectrometry modalities. GC-MS is an efficient technique for detecting metabolites with mass weights typically < 500 Dalton, providing it with the ability to readily detect key compounds of primary metabolism including small organic acids and amino acids, and differentiate between sugar isomers (e.g., glucose, fructose, mannose, sorbose, etc.) (Smith and Morowitz, 2004). We detected 55 metabolites of these types with the GC-MS technique compared to the 16-20 metabolites detected by the MALDI-MSI technique. On the other hand, the MALDI-MSI capability used in this work is a powerful technique for detecting aromatic compounds and secondary metabolites. We detected about 73 metabolites of these types using MALDI-MSI compared to the 25 metabolites identified by the GC-MS approach (supplementary tables S2 and S3). Another possible explanation for the discrepancy between MALDI-MSI and GC-MS results is that since with MALDI-MSI we capture only a 10µm plane of the whole leaf, there is a chance that other areas of the leaf (within palisade and vascular regions) show a different trend of metabolite accumulations which in average could resemble the GC-MS output. However, this point is at least partially mitigated by a large number of biological and technical replicates used in this study for generating metabolic snapshots by MALDI-MSI. Moreover, a recent remarkable development in the MALDI-MSI approach, which uses a novel on-tissue chemical derivatization strategy (Zemaitis et al., 2023), has significantly expanded the metabolite coverage of phytocompounds including primary carbohydrates, amino acids, and TCA cycle compounds. This updated version of MALDI-MSI can be adopted in future studies to improve the cross-comparison between the whole tissue and spatial metabolomics techniques.

Overall, using spatial metabolomics, we found novel distinct WD-responsive metabolic maps of two major leaf cell types in poplar, which provide a better understanding on how different cell types work together to respond to environmental perturbations in a strategic perennial tree. Understanding spatiotemporal differences in metabolic responses to WD stress will help engineer targeted pathways in specific cell types, potentially achieving maximum WD response and avoiding any pleiotropic effects on plant growth and yield.

### Data availability statement

The datasets generated and analyzed during the current study will be made available in the respective data repositories shortly after submission. All the MALDI-MSI datasets and annotations will be publicly available at METASPACE. METASPACE links for reviewers:

- Positive ion mode analysis: https://metaspace2020.eu/api_auth/review?prj=e73ebe52-3b62-11ed-89bf-536c64910508&token=K3GB4ZCy0yDe
- Negative ion mode analysis: https://metaspace2020.eu/api_auth/review?prj=60c05570-3b80-11ed-89bf-0f161aeeebf3&token=sD4R5421iN8E

## Author contributions

**Vimal Kumar Balasubramanian** designed and performed experiments, contributed to data analyses, wrote the original draft, reviewed, and edited the manuscript; **Dusan Velickovic** performed MALDI-MSI metabolomics and performed data analyses, reviewed, and edited the manuscript; **Maria Del Mar Rubio Wilhelmi** performed poplar water-deficit treatments and generated biomass data and helped generating GC-MS based whole tissue metabolomics data, reviewed, and edited the manuscript; **Christopher Anderton** assisted with the MALDI-MSI workflow development and data analysis, reviewed, and edited the manuscript; **Neal Stewart, Jr., Stephen DiFazio, and Eduardo Blumwald** acquired funding, reviewed, and edited the manuscript; **Amir H. Ahkami**: conceptualized the work, acquired funding, designed experiments, wrote and edited the manuscript.

## Funding

This work was supported by funding from the Biological and Environmental Research (BER) in the U.S. Department of Energy (DOE) Office of Science, Genomic Science Program, Biosystems Design to Enable Next-Generation Biofuels (SyPro Poplar project, Award Number: DE-SC0018347). Part of this work was conducted at EMSL (Environmnetal Molecular Sciences Laboratory), a DOE Office of Science User Facility sponsored by the Office of Biological and Environmental Research and operated under Contract No. DE-AC05-76RL01830, located at Pacific Northwest National Laboratory (PNNL).

## Acknowledgments

We thank Dr. Albert Rivas Ubach (National Institute for Agricultural and Food Research and Technology (INIA-CSIC) for his help in one-way ANOVA analysis of the GC-MS metabolomics dataset.

## Conflict of interest

The authors declare no conflicts of interest.

## Supplementary material

**Supplementary Table S1:** Shoot and root biomass and gas exchange parameters (photosynthesis (A), conductance (gs) and transpiration (E)) measured from early water deficit (E-WD) and late water deficit (L-WD) stressed and recovered (R) poplar leaf tissue. Data averaged from four biological replicates for biomass analysis and six biological replicates for leaf gas exchange parameters. t-test (p<0.05) was used for statistical analysis.

**Supplementary Table S2:** Total number of significantly altered metabolites identified through MALDI-MSI analysis of early water deficit (E-WD) and late water deficit (L-WD) stressed and recovered (R) poplar leaf tissue. Positive and negative ionization modes were used for data acquisition and non-overlapping metabolic features were merged. Metabolites were grouped into various metabolic classes by Van Krevelen classification, and cell-type unique (palisade unique and vascular unique) metabolites were separately tabulated from cell-type shared metabolites. Fold change values were calculated based on relative abundance of each metabolite in stress vs control conditions. T-test analysis was used for statistical analysis. Metabolites with significant (n=97, p<0.05) and moderate significant (n=7, 0.05<p<0.1) changes are included.

**Supplementary Table S3:** Total number of significantly altered metabolites identified by whole leaf metabolite profiling using GCMS technique from early water deficit (E-WD) and late water deficit (L-WD) stressed and recovered (R) poplar leaf tissue. Fold change values were calculated based on relative abundance of each metabolite in stress vs control conditions. T-test analysis was used for statistical analysis (p<0.05).

**Supplementary Table S4:** Total number of significantly altered metabolites identified through negative ionization mode analysis in MALDI-MSI from early water deficit (E-WD) and late water deficit (L-WD) stressed and recovered (R) poplar leaf tissue. Fold change values were calculated based on relative abundance of each metabolite in stress vs control conditions. T-test analysis was used for statistical analysis and the data from this table is used to created Sup. Table S2.

**Supplementary Table S5:** Total number of significantly altered metabolites identified through positive mode analysis in MALDI-MSI from early water deficit (E-WD) and late water deficit (L-WD) stressed and recovered (R) poplar leaf tissue. Fold change values were calculated based on relative abundance of each metabolite in stress vs control conditions. T-test analysis was used for statistical analysis and the data from this table is used to created Sup. Table S2.

---

**Supplementary Figure S1.**
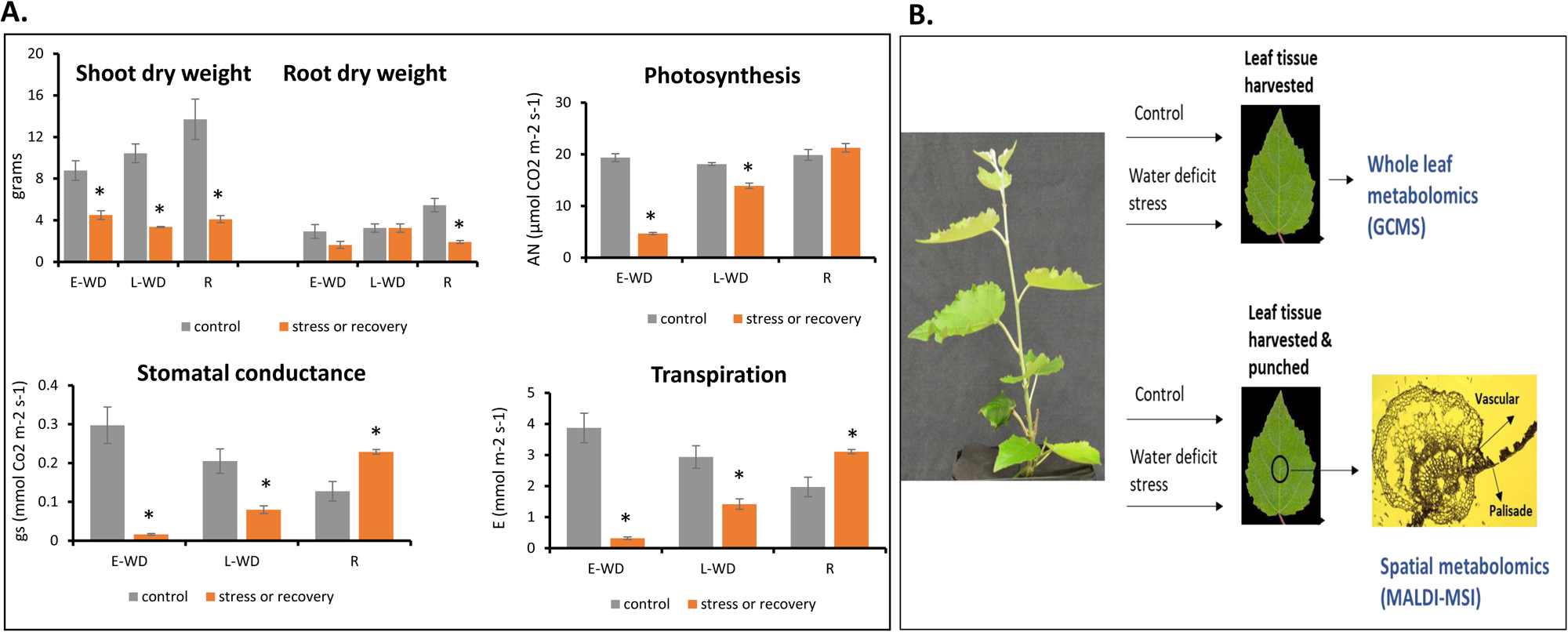
Water deficit stress alters plant dry weight and leaf physiological parameters in poplar. **(A.)** Shoot and root biomass and gas exchange parameters, photosynthesis (A), conductance (gs) and transpiration (E) measured from plants exposed to early water deficit (E-WD) stress (30-35% relative SWC), late water deficit (L-WD) stress (water level maintained for 10d at 30-35% relative SWC) and recovery from stress (R) and data were collected. Data averaged from four biological replicates and t-test was used for statistical analysis. *represents pvalue<0.05. (**B.)** Leaf samples were collected for whole leaf tissue and spatial cell type specific metabolite profiling. Six biological replicates were harvested in any given condition. Matrix-assisted laser desorption Ionization-mass spectrometry imaging (MALDI-MSI) and Gas chromatography-mass spectrometry (GC-MS) analysis.

**Supplementary Figure S2.**
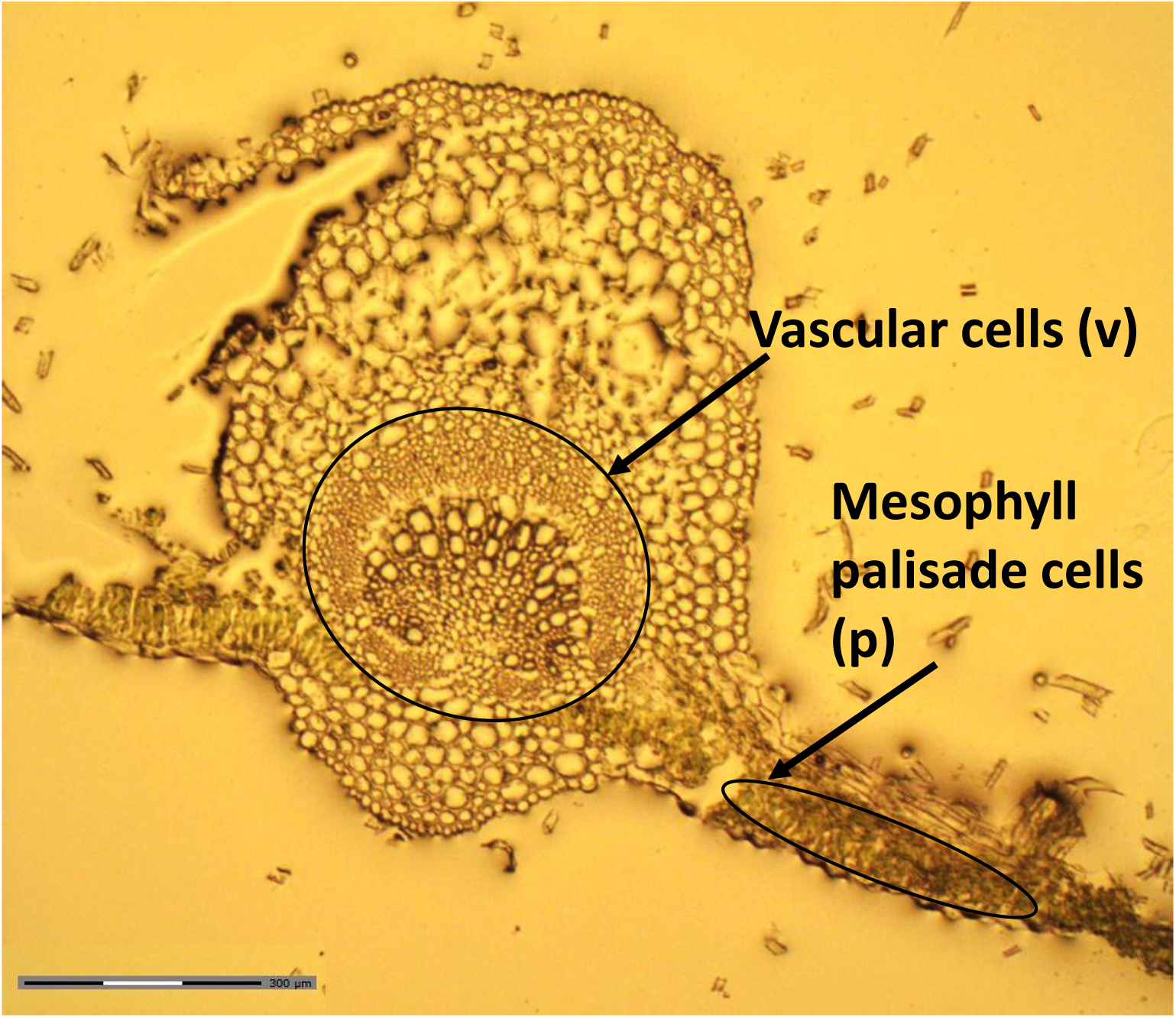
Cryosectioning of a poplar leaf tissue. Cryosection highlighting mesophyll palisade (p) and vascular (v) cell types used for MALDI MSI to generate spatial metabolome data. Scale bar represents 300µm.

**Supplementary Figure S3.**
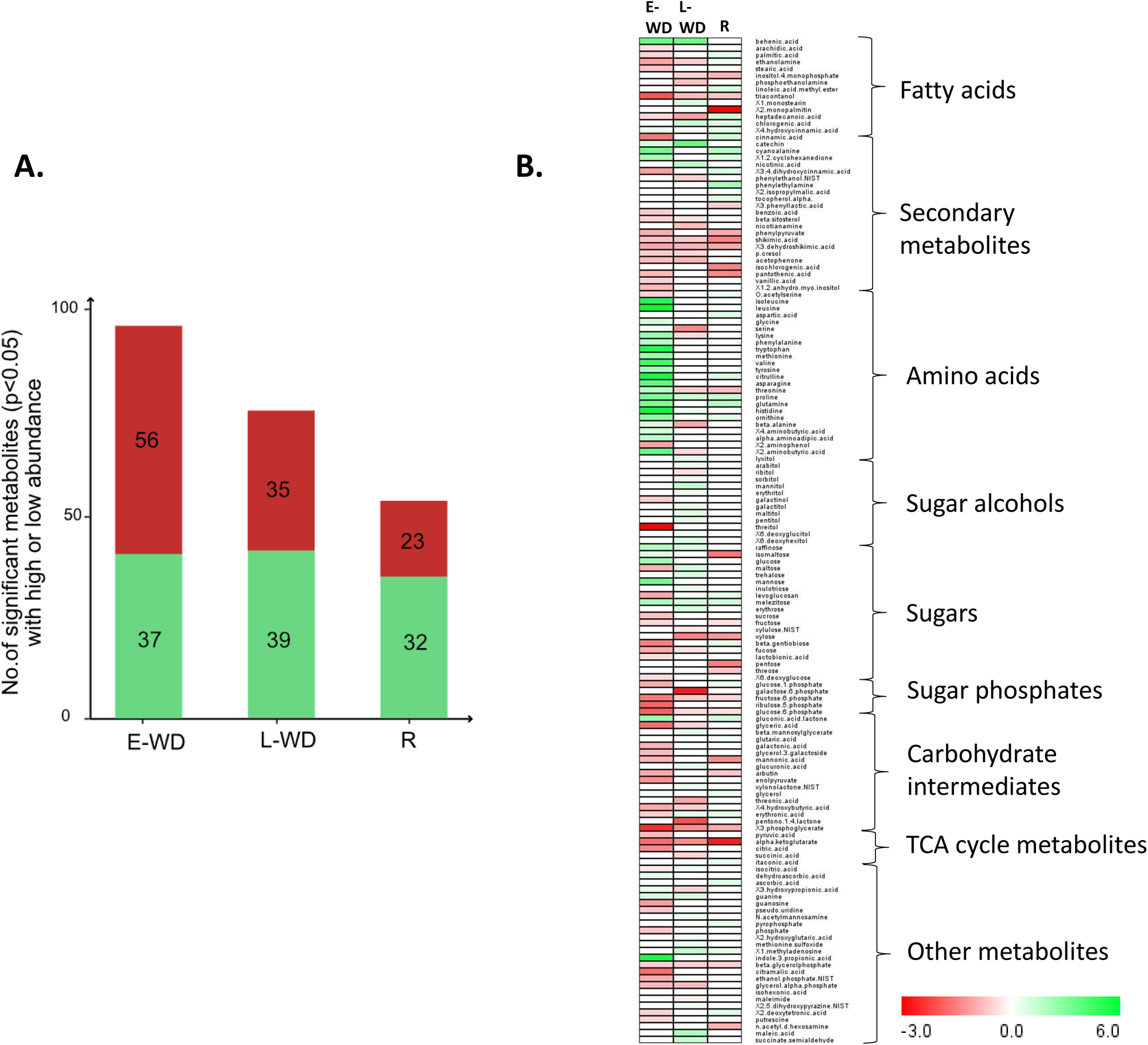
Whole leaf tissue-based metabolomics identified water deficit stress responsive metabolites. **(A.)** Total no. of metabolites significantly altered during early (E-WD) and late (L-WD) water deficit stress and recovery (R) conditions in whole leaf tissue of poplar. Green and red bars highlight metabolites with increased or reduced abundance levels during stress and recovery conditions. Data averaged from three biological replicates and one-way ANOVA was used for statistical analysis (p<0.05). (**B.)** Total significant metabolites were categorized into broader metabolic classes using KEGG pathway analysis. The heatmap shows metabolite abundance levels in E-WD, L-WD and R conditions. Data averaged from 3 biological replicates and ANOVA (p<0.05) was used for statistical analysis.

## Notes

### Competing Interest Statement

The authors have declared no competing interest.

